# Mechanistic learning to predict and understand minimal residual disease

**DOI:** 10.64898/2026.04.16.718968

**Authors:** Sadegh Marzban, Mark Robertson-Tessi, Jeffrey West

## Abstract

Mechanistic modeling has long been used as a tool to describe the dynamics of biological systems, especially cancer in response to treatment. Their key advantage lies in interpretability of relationships between input parameters and outcomes of interest. Mechanistic models may also be calibrated to a cohort of patients and scaled up to generate a simulated set of virtual patients whose aggregate behavior reproduces key characteristics of the real patient population. In contrast, machine learning techniques offer strong prediction performance, especially for high dimensional datasets that are common in oncology. Here, we employ a Mechanstic Learning framework that combines the advantages of both approaches by training machine learning models on mechanistic parameters inferred from clinical patient data. We assess the ability of virtual clinical cohorts for the purpose of 1) scaling up small cohort sizes and 2) balancing unbalanced patient subgroups in the setting of BCR::ABL1 positive lymphoblastic leukemia. Our mechanistic model (a Markov chain model) contains sixteen parameters that describe the rate of cell fate transitions that occur in patients with B-cell precursor acute lymphoblastic leukemia. The machine learning (a ridge logistic regression model) is trained on these parameters to predict two clinically-relevant features: BCR::ABL1 fusion gene status (positive or negative) and minimal residual disease status (positive or negative) post-induction chemotherapy. Model training is done in an iterative fashion to assess which (and how many) parameters are critical to maintain high predictive performance. Using machine learning models trained on the clinical flow-cytometry data, we find that the stem-like cell state alone is the most predictive feature for both BCR::ABL1-positive and MRD-positive disease, with composite scores (defined as the average of accuracy, balanced accuracy, and area under the curve) of 0.80 and 0.67, respectively. By comparison, mechanistic learning achieves comparable or improved composite scores for BCR::ABL1-positive and MRD-positive disease, with scores of 0.81 and 0.71, respectively, using only de-differentiation for BCR::ABL1 and stem-state persistence together with differentiation-directed exit for MRD. Virtual Patient (VP) expansion is informative for robustness analysis and class balancing, but full cohort expansion introduced additional heterogeneity, reduced predictive performance, and required larger models, whereas VP-based balancing yielded only a modest gain over class weighting at substantially greater computational cost. In summary, a mechanistic-learning approach not only preserves predictive performance, but also provides a biological hypothesis for why stemness is predictive of these clinically relevant outcomes.

## 1 Introduction

The cancer stem cell (CSC) hypothesis proposes that tumor growth and therapeutic resistance can be driven by a relatively small subpopulation of self-renewing cells[1]. In acute myeloid leukemia (AML), this idea was supported by transplantation studies showing that leukemia-initiating activity was enriched in the CD34^+^CD38^−^ compartment[2], and by subsequent work showing hierarchical organization from primitive hematopoietic-like cells to more differentiated leukemic populations[3]. These primitive AML cells also express high levels of ATP-binding cassette transporters, consistent with multidrug resistance and long-term persistence under therapy[4]. Clinically, higher frequencies of CD34^+^CD38^−^ cells in AML have been associated with higher MRD, worse survival, and greater relapse risk[5–8].

In B-cell acute lymphoblastic leukemia (B-ALL), however, the role of stem-cells is less rigid. Leukemia-initiating potential is not confined to a single CD34/CD38-defined compartment: both CD34^+^CD38^+^CD19^+^and CD34^+^CD38^−^CD19^+^populations can propagate disease[9], MLL-rearranged ALL can initiate from both CD34^+^and CD34^−^ fractions[10], and CD34/CD38 expression itself can be reversible rather than strictly hierarchical[11]. Even so, immature CD34^+^CD38^−^ populations remain clinically important, because their abundance at diagnosis is associated with higher MRD and poor prognosis in childhood B-ALL[12–14]. Together, these observations suggest that the key biological problem in B-ALL is not simply the presence of a fixed stem-cell compartment, but the dynamic movement of cells among stem-like, proliferative, and quiescent phenotypic states.

### 1.1 Mechanistic modeling in ALL

Mechanistic modeling provides a principled way to represent how biological systems evolve over time[15, 16] and has been widely used to study proliferation, differentiation, treatment response, and resistance in cancer[17–23]. Its main advantage is interpretability: model parameters correspond to biological processes, so the fitted model can help explain not only what predicts outcome, but also why. In the present setting, Markov-state models are especially attractive because they convert static flow-cytometry snapshots into inferred transition tendencies among phenotypic states[15, 24]. In our previous work, we used this framework to estimate patient-specific transitions among four CD34/CD38-defined B-ALL states: CD34^+^/CD38^−^, CD34^+^/CD38^+^, CD34^−^/CD38^+^, and CD34^−^*/*CD38^−^[24]. That analysis suggested that poor-prognosis patients are characterized by enhanced stem-like self-renewal and increased inward flux toward the stem-like compartment. However, mechanistic models alone can be limited by the simple mathematical form of their underlying assumptions and they are typically used explain the behavior of a single patient in isolation[25]. This creates a natural tension between mechanistic interpretability and predictive accuracy.

### 1.2 Machine learning models of ALL outcomes

Data-driven machine learning addresses the problem of maximizing predictive performance from high-dimensional data[26–30]. In acute leukemia, machine-learning methods have already been applied to relapse prediction in childhood ALL, mortality and relapse stratification in pediatric ALL, survival modeling in Philadelphia chromosome-like ALL, short-term mortality prediction in AML, genomic risk subclassification in AML, and machine-learning-guided MRD analysis in AML[31–36]. Across oncology more broadly, supervised learning approaches have proven useful for risk stratification, biomarker discovery, and clinical outcome prediction from clinical, molecular, and imaging variables[37–40]. Yet purely data-driven models often provide limited biological explanation and interpretability, even when they perform well. This is a particular problem in B-ALL, where we would like to know not only which phenotype predicts BCR::ABL1 status or MRD, but how that prediction reflects the dynamic transitions of cells between stem-like and differentiated phenotypic states.

### 1.3 Mechanistic models to generate virtual patients

Machine learning algorithms are typically data-intensive, leading to various proposed methods for scaling up patient cohort sizes. For example, virtual clinical cohorts are simulated patient populations constructed so that their aggregate behavior reproduces key characteristics of a real patient population, while remaining grounded in mechanistic or systems-pharmacology models[41– 43]. Related patient-specific digital twin and virtual cohort approaches have also been developed to forecast treatment-associated toxicity and support treatment decision-making[44–46]. In oncology, these virtual cohorts can be used to test a wide range of treatment schedules, treatment options, and sequential ordering of therapies in a safe and cost-effective computational framework before prospective evaluation in patients[42, 43]. An additional practical advantage is that virtual cohorts can facilitate data sharing while protecting patient privacy, because they can be generated to reflect cohort-level properties without exposing patient-level records directly[42]. At the same time, their utility depends on model structure, calibration quality, and whether the simulated heterogeneity adequately captures real patient variability; thus, virtual cohorts should be viewed as a complement to, rather than a replacement for, clinical data[41, 42]. In this manuscript, we explore the utility of virtual cohorts in scaling up cohort sizes in small datasets and to balance datasets with an unbalanced number of samples in each subgroup (e.g., below we have many more samples from patients who are negative for minimal residual disease than positive).

### 1.4 Mechanistic learning combines predictive power with interpretability

Mechanistic learning is designed to bridge the gap between mechanistic interpretability and predictive accuracy by combining mechanistically derived features with machine-learning classifiers[47–51]. In this framework, machine learning provides discrimination and generalization, whereas the mechanistic representation preserves biological meaning. Here, we use a sequential mechanistic-learning strategy[48]: we first infer patient-specific transition structure among the four CD34/CD38 states, then use those derived quantities as predictors for downstream classification tasks. This allows us to test whether a compact set of transition features can match or improve upon flow-cytometry-based prediction while also explaining the transition programs associated with genotype and treatment response.

Mechanistic learning is therefore not simply a model-ensemble strategy; it is a way to turn interpretable disease dynamics into predictive variables. We apply our Mechanistic Learning framework to predict two important clinical categories: the status of the BCR::ABL1 fusion gene (positive or negative, with BCR::ABL1-like patients excluded) and the status of minimal residual disease (MRD) post-inducation chemotherapy. We assess the role of BCR::ABL1 because it is a known driver of worse outcomes in ALL[52], and we assess post-induction chemotherapy MRD because this determines whether patients receive consolidation or intensification chemo after induction chemotherapy. Below, we show that the development of accurate predictive models are possible in both cases, but that the number of biological or mechanistic features needed for accurate predictions can be drastically reduced, leading to an increased interpretability of the driver of disease.

Our working hypothesis is that machine learning classifiers trained on mechanistically derived transition features will retain competitive predictive accuracy while offering clearer biological explanations of prognosis-related phenotypes[47, 48]. In particular, we hypothesize that BCR::ABL1 status will be associated with transition patterns that promote re-entry into the stem-like compartment, whereas MRD will be associated more strongly with persistence of that compartment once established.

## 2 Methods

As summarized in Figure 1, we implement a sequential mechanistic-learning pipeline that begins with clinical flow-cytometry features (Step 0 in Figure 1) and then uses the mechanistic modeling parameters (Step 1) parameterized on patient cohorts (Step 2) for downstream prediction algorithms (Step 3). Finally, we iteratively reduce the number of mechanistic features fed to machine learning algorithms (Step 4) to identify a minimal feature set, for increased interpretability (Step 5).

**Figure 1:**
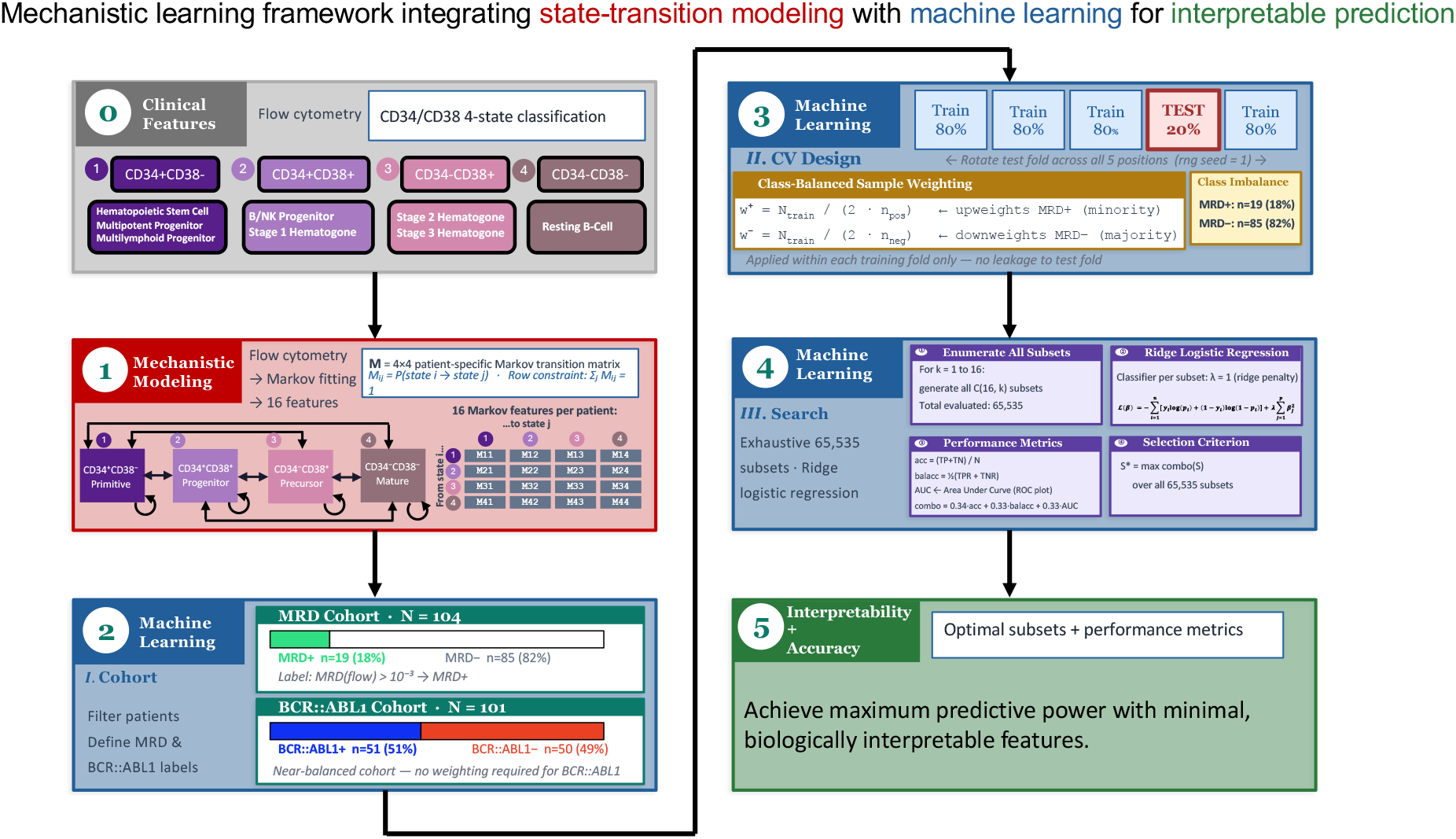
Mechanistic-learning pipeline from flow cytometry to interpretable prediction of clinical outcomes. Schematic overview of the framework integrating mechanistic modeling with machine learning. (1) Mechanistic feature construction: multiparametric flow-cytometry data define four CD34/CD38 cell states, from which patient-specific discrete-time Markov transition matrices are inferred to generate mechanistically grounded features (16 transition rates per patient). (2) Cohort construction: patients are stratified by measurable residual disease (MRD) and BCR::ABL1 status to form labeled datasets for supervised learning. (3) Class-imbalance handling: weighted learning addresses skewed class distributions, particularly for MRD classification. (4) Model training and feature selection: exhaustive subset search with ridge logistic regression identifies low-dimensional feature sets that optimize predictive performance (accuracy, balanced accuracy, AUC, and combination score) under cross-validation. (5) Interpretation and prediction: the framework achieves strong predictive performance with a minimal set of biologically interpretable features, linking transition dynamics such as self-renewal and differentiation to clinical outcomes.

### 2.1 Description of the clinical dataset

Figure 2 summarizes the clinical setting of this study. Patients follow a typical B-ALL treatment course with serial evaluation of MRD and relapse-related outcomes (Figure 2A). This cohort combines cytogenetic and sequencing information with matched flow-cytometry measurements of the four CD34/CD38-defined states (Figure 2B,C). Patients are stratified by BCR::ABL1 status, a clinically important molecular subtype[53], and the dataset also includes remission samples for comparison. For each patient, we also have previously inferred state-transition rates among the four CD34/CD38 compartments[24], summarized as patient-level averaged Markov transition rates (see below) for 100 stochastic training runs and shown schematically in Figure 2D. These are referred to as our dataset’s mechanistic features, since they represent the parameters from our mechanistic model, a Markov chain model. Even before formal prediction, these mechanistic features reveal biological organization. Hierarchical clustering highlights transitions into the CD34^+^/CD38^−^ state as a major axis of variation, particularly in BCR::ABL1-positive disease. Figure 3 further suggests that MRD-positive and BCR::ABL1-positive patients share a transition pattern characterized by enhanced inward flux toward the stem-like compartment, whereas MRD-negative and BCR::ABL1-negative patients show weaker reconstruction of that state. These visual patterns motivate the central question of this study: can mechanistically inferred transition features provide predictive accuracy comparable to or better than standard flow-cytometry features, while simultaneously explaining why specific phenotypic states and transition programs are associated with genotype, disease persistence, and treatment effectiveness?

**Figure 2:**
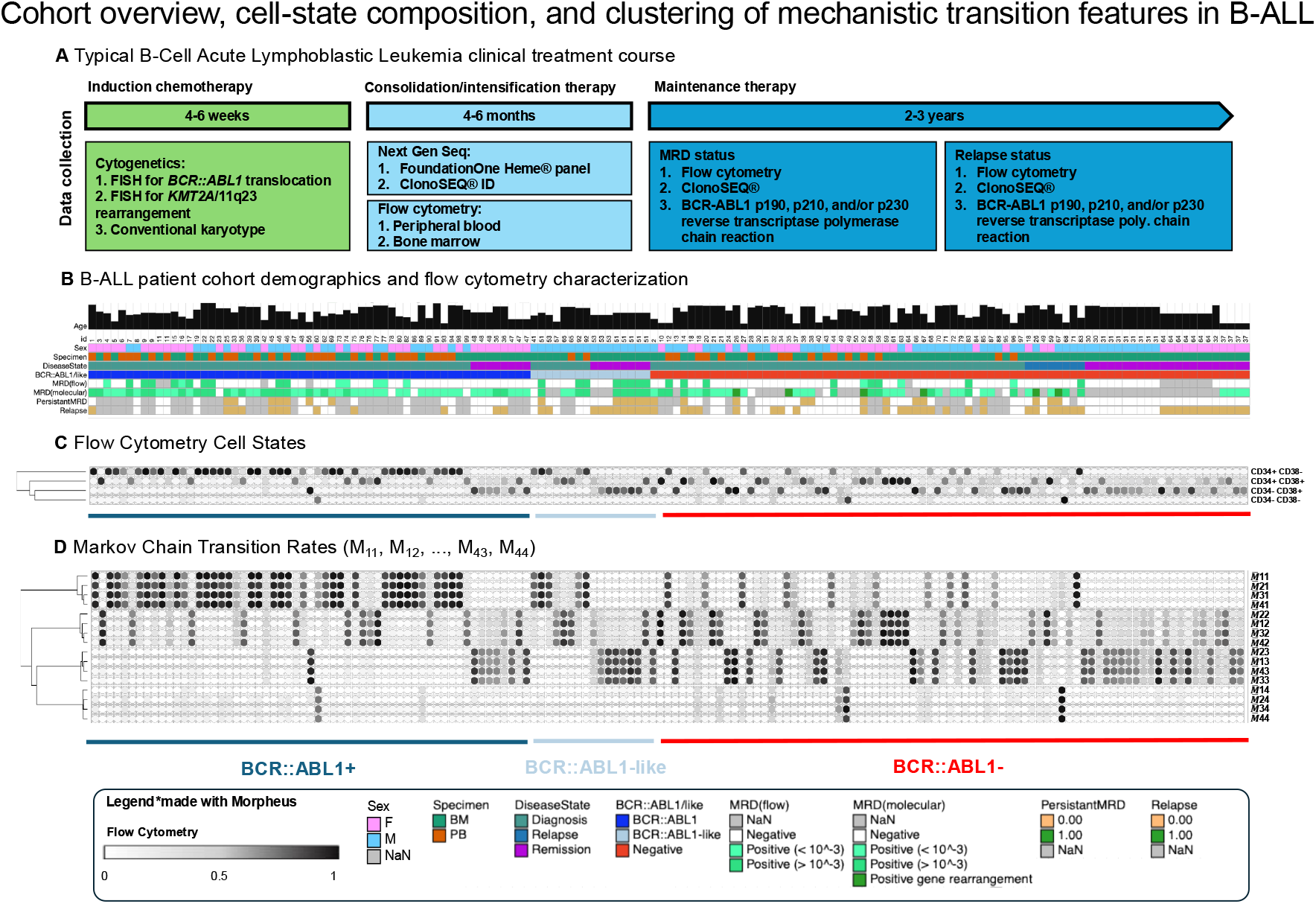
Cohort overview, cell-state composition, and clustering of mechanistic tran-sition features in B-ALL. (A) Schematic of the clinical treatment course for B-cell acute lymphoblastic leukemia (B-ALL), including induction, consolidation, and maintenance phases, along with associated data collection modalities (cytogenetics, next-generation sequencing, and flow cytometry).(B) Patient cohort overview with clinical annotations, including disease state, specimen type, measurable residual disease (MRD), and BCR::ABL1 status.(C) Flow cytometry-derived cell-state composition (X features), where each patient is represented by the relative abundance of four CD34/CD38-defined immunophenotypic states.(D) Patient-level averaged Markov transition rates 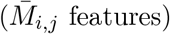, calculated as the mean over 100 stochastic training runs for each patient and capturing transition dynamics between cell states. Hierarchical clustering of transition features reveals structured patterns across patients, with distinct clustering of transition profiles associated with BCR::ABL1 status. In particular, patients with BCR::ABL1-positive disease exhibit coherent transition patterns indicative of enhanced self-renewal and inward flux toward stem-like states, while BCR::ABL1-negative patients display more heterogeneous differentiation.

**Figure 3:**
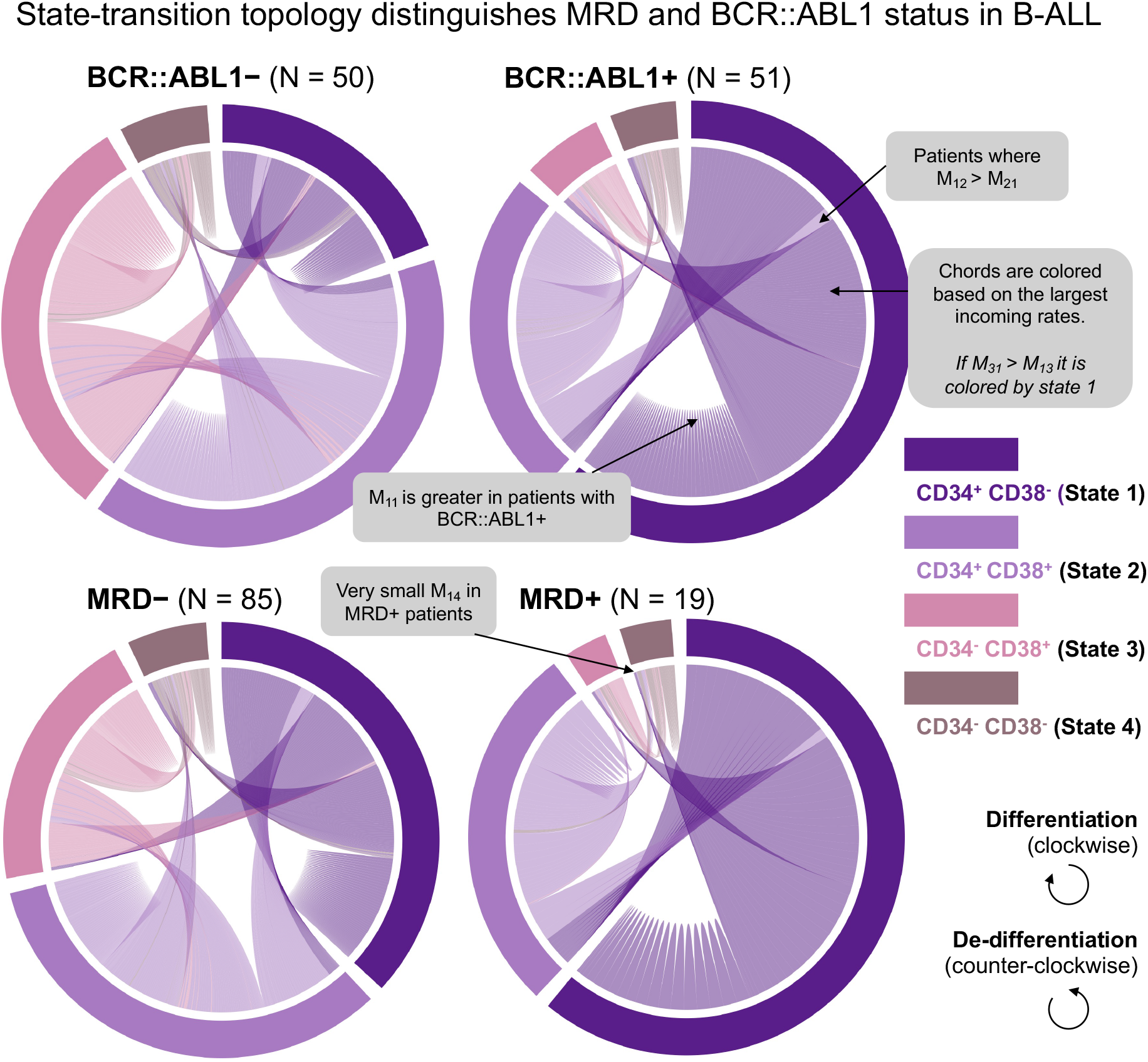
Transition dynamics across immunophenotypic states stratified by BCR::ABL1 and MRD status. Chord diagrams summarize patient-level averaged inferred transition rates, calculated as means over 100 stochastic training runs, between four CD34/CD38-defined states (State 1: CD34^+^/CD38^−^; State 2: CD34^+^/CD38^+^; State 3: CD34^−^/CD38^+^; State 4: CD34^−^/CD38^−^) for individual patients grouped by BCR::ABL1 status (top) and measurable residual disease (MRD) status (bottom). Chord widths represent transition magnitude, and colors denote the dominant incoming state for each transition. Across patients, MRD+ and BCR::ABL1-positive groups exhibit qualitatively similar transition structures, characterized by enhanced flux toward the stem-like CD34^+^/CD38^−^ compartment (State 1). Self-renewal and inward transitions into State 1 are elevated in MRD+ and BCR::ABL1-positive patients, indicating active reconstruction of stem-like populations.

### 2.2 Mechanistic model: Markov chain

As a point of departure, we take the mechanistic modeling framework developed and parameterized in Gravenmier et. al. [24], which is a Markov chain transition matrix that quantifies the rate of transition from cell type *i* to cell type *j* as *M*_*i,j*_. Four cell types are considered (CD34^+^/CD38^−^, CD34^+^/CD38^+^, CD34^−^/CD38^+^, CD34^−^/CD38^−^) as shown in Figure 2C. We term these our clinical features:

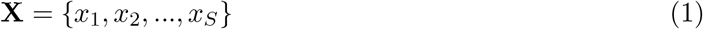

In our dataset, we have *S* = 4 clinical features (Figure 2C). To estimate the transition dynamics among these four states, we used a mechanistic model, here a Markov chain model, with the following set of parameters:

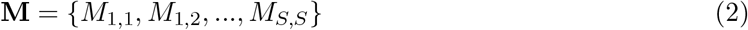

with, by definition, *S*^2^ mechanistic features. A Markov transition matrix, **M**, has an associated steady-state given by the following stationary equation:

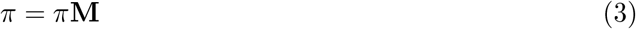

In our previous work[24], we determined the set of Markov transition rate parameters, **M**, which minimize the least square error between the steady-state *π* and each patient’s biological data, **X** such that the mechanistic model describes the data within a reasonably low error, *ϵ*

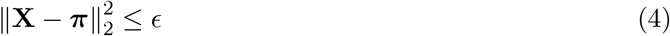

### 2.3 Defining a set of virtual patients

As stated above, virtual cohorts are often used to scale or balance datasets. Here, we construct virtual patients by repeating the mechanistic model fitting process *N* times to generate *N* virtual patients. That fitting process begins with an initial guess of the Markov transition matrix, **M**, and iterates according to the Newton–Mason algorithm[15] until the associated steady-state *π* converges to the target, **X**. The Newton–Mason training algorithm is stochastic, and therefore each training run yields a unique Markov transition matrix that satisfies eqn. 4. We repeat the training process *N* = 100 times, and we refer to the resulting matrices **M**^1^, **M**^2^, …, **M**^*N*^ as “virtual patients,” such that each real patient dataset **X** has *N* associated virtual-patient (VP) parameter sets. For analyses that require a single mechanistic representation per real patient, we define each transition feature as the average over that patient’s *N* virtual realizations. Specifically, if 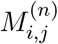 denotes the transition parameter from state *i* to state *j* from the *n*th virtual patient, then we use

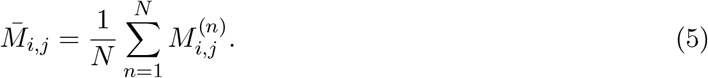

These values are visualized in figure 3 in a chord diagram, repeated across BCR::ABL1 positive/negative and MRD positive/negative subgroups. Visually, there are some notable differences in cell state transitions across subgroups, such as a higher stem state self-renewal, *M*_11_, as previously reported[24]. Rather than rely on visual inspection, we employ machine learning algorithms to identify key mechanistic features that distinguish these clinical subgroups.

### 2.4 Exhaustive subset search with ridge logistic regression

The endpoint datasets were class-imbalanced (Step 2 in Figure 1): MRD classification used *N* = 104 samples (*n* = 85 for MRD-negative, *n* = 19 for MRD-positive), and BCR::ABL1 classification used *N* = 101 samples (*n* = 50 for BCR::ABL1-negative, *n* = 51 for BCR::ABL1-positive). To address class imbalances (step 3 in Figure 1), we used class-balanced weighting within each stratified 5-fold training split (i.e., using only the 4/5 training partition in each fold). Let *N*_train_ = (4*/*5)*N* and *n*_train_ = (4*/*5)*n* for each of the two classes (positive vs negative). Each training sample received weight

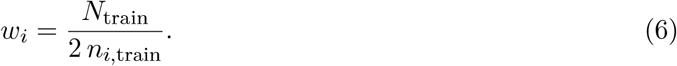

Accordingly, MRD fold weights were 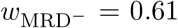 and 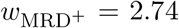. For each endpoint (MRD status and BCR::ABL1 status), we performed an exhaustive subset search as part of model training and feature selection (Step 4 in Figure 1). For mechanistic predictors, we searched over the 16 Markov transition features; for each subset size *k* = 1, …, 16, we enumerated all 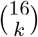 possible feature combinations (total 2^16^ − 1 = 65,535 non-empty subsets). For comparison of predictive power, we also fed the clinical flow-cytometry features (Step 0 in Figure 1) directly into the same machine-learning pipeline; with four clinical features (*x*_1_, *x*_2_, *x*_3_, *x*_4_), the exhaustive search used 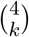 candidate subsets. Each candidate subset was evaluated using weighted ridge logistic regression under stratified 5-fold cross-validation. We chose ridge regularization because the Markov transition-rate predictors are intrinsically correlated: each transition matrix is row-stochastic (i.e., probabilities sum to 1), which induces within-row dependence among transition rates. In logistic regression, ridge penalization is well suited to this setting because it stabilizes coefficient estimates under multicollinearity and can improve predictive performance through coefficient shrinkage[54, 55].

Given feature vector **x**_*i*_ and label *y*_*i*_ ∈ {0, 1}, the fitted probability was

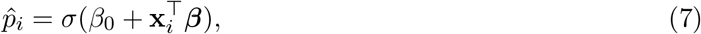

where *σ*(*z*) = 1*/*(1 + *e*^*−z*^). Parameters were estimated by minimizing the weighted penalized negative log-likelihood

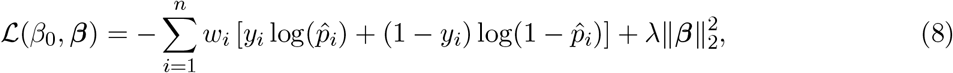

with class weights *w*_*i*_ and ridge penalty *λ* = 1.

### 2.5 Performance metrics and information criteria

Next, we report both accuracy metrics and information criteria for each predictive model. This approach allows us to maximize model accuracy, while minimizing the number of parameters given to the classifier. By reducing the number of parameters, the interpretability of the model is increased. For each subset, we recorded cross-validated accuracy, balanced accuracy (BalAcc), and Area Under the Curve (AUC), consistent with the model-selection workflow in Figure 1 (step 4). To rank subsets by the combination of these predictive metrics, we defined a composite score

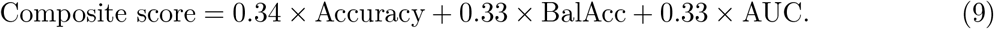

Model selection used this composite score together with information criteria to favor both predictive performance and parsimony. Let *n* denote sample size and ℓ the maximized log-likelihood. Because ridge regularization shrinks coefficients, we used the effective degrees of freedom

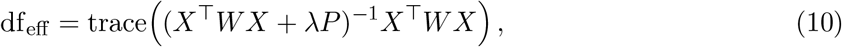

where *W* is the logistic working-weight matrix and *P* is the ridge penalty matrix. We then computed Bayesian Information Criterion (BIC) [56].

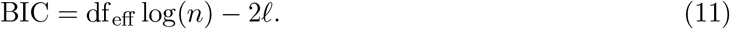

We employed BIC to guide feature selection, as its stronger complexity penalty (log *n*) favors parsimonious models [56, 57]. Given the exhaustive subset search and the goal of identifying stable, mechanistically interpretable features, BIC provides a principled safeguard against overfitting while favoring models with stronger empirical support. We opt for the set of parameters which minimizes BIC, but it is conceivable that some clinical applications may prefer a high accuracy over and above high interpretability (or vice versa).

### 2.6 Virtual-patient augmentation and resampling analyses

We used virtual-patient (VP) realizations in two secondary analyses. First, to assess whether VP-based data augmentation improved prediction, we treated the full set of 100 VP parameterizations associated with each real patient as separate mechanistic observations, assigned each VP the clinical label of its originating real patient, and reran the same weighted ridge-logistic subset-search pipeline used in the primary analyses. To avoid information leakage, the five cross-validation folds were assigned at the level of the originating real-patient ID, so no VP realizations from the same patient appeared in both the training and test sets within any fold. This produced a 100-fold expanded mechanistic cohort for BCR::ABL1 and MRD prediction.

Second, as a complementary sensitivity analysis for class imbalance, we constructed balanced VP datasets by repeated resampling from the *N*_VP_ = 100 realizations associated with each real patient until the smaller subgroup has an equivalent number of samples as the larger. For example, the MRD positive subgroup is four to five times smaller, and therefore we must select four or five VPs from each MRD positive patient so that the expected class sizes were approximately balanced. We then reran the same subset-search pipeline on each resampled dataset and summarized the resulting feature-selection patterns across repetitions.

To avoid cherry-picking and ensure broad coverage when sampling from the VP cohort, we calculated the minimum number of resampling repetitions needed so that each VP realization would have high probability of being included at least once across the analysis. Let *v* denote the number of virtual patients per real patient desired to balance the dataset. For example we select *v* = 85*/*19 for the MRD positive subgroup. Under uniform within-patient sampling, the probability that a specific VP realization is selected after *B* fold repetitions is

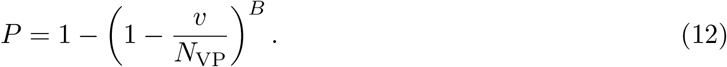

If we require each VP realization to have probability at least *P* = 0.95 of being sampled at least once, then the number of repetitions must satisfy equation 12 solved for *B*:

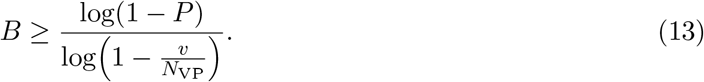

For MRD negative patients, *v*_−_ = 1, so achieving *P* = 0.95 gives *B* ≥≈298. For MRD positive patients, balancing to 85 negative samples per repetition corresponds to an average draw size of *v*_+_ = 85*/*19 ≈4.47 VPs per patient, which yields *B ≥* 65.5. Thus, the MRD negative class is the limiting case, and we used the corresponding minimum number of balanced VP resampling repetitions, namely *B* = 300, so that each MRD negative VP realization had approximately a 95% probability of being included at least once across the analysis.

## 3 Results

### 3.1 Real-cohort flow-cytometry and averaged Markov features

We sought to predict two relevant clinical categories: the status of each patient for BCR::ABL1 fusion gene (positive or negative, with BCR::ABL1-like patients excluded) and the status of minimal residual disease post-inducation chemotherapy. Applying the workflow described in Methods (Figure 1), we ranked all Markov-feature subsets using cross-validated predictive metrics and information criteria (Figure 4). We first apply the machine-learning pipeline to baseline flow-cytometry features (Figure 4A; Supplementary Table S1). Among flow-cytometry subsets for BCR::ABL1 prediction, the best clinical feature was the single-state feature *x*_1_ (state 1, stemness-associated compartment), which achieved composite score 0.80 and BIC 101.47.

**Figure 4:**
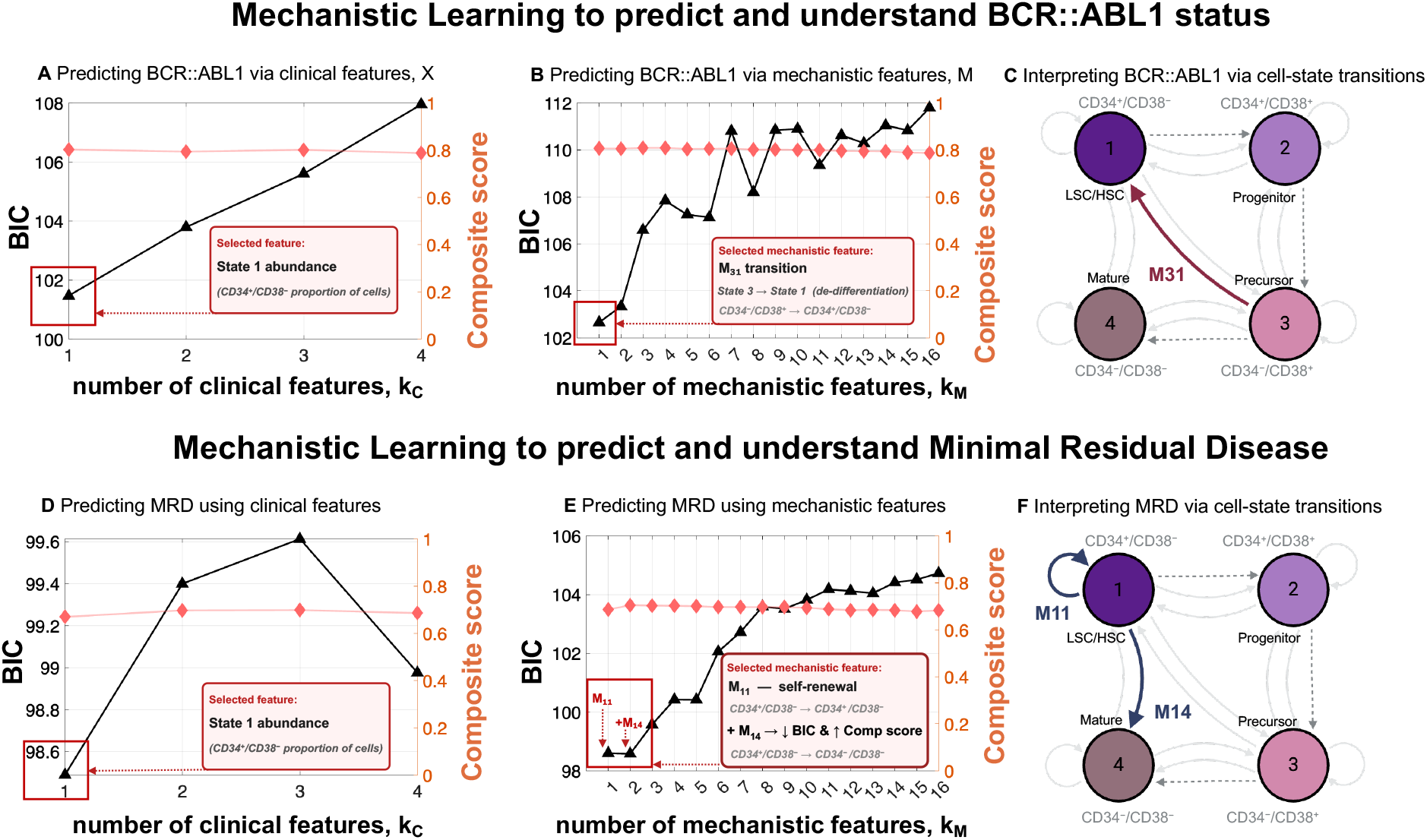
Mechanistic features enable accurate and interpretable prediction of genotype and clinical outcome. Top row: predicting BCR::ABL1 and interpreting mechanistic drivers. Bottom row: predicting measurable residual disease (MRD) and interpreting mechanistic drivers. (A–B, D–E) Model selection comparing clinical and mechanistic features. Model fit (BIC) and predictive performance (composite score) are shown as a function of feature subset size. Mechanistic features achieve comparable or improved performance. (C, F) Mechanistic interpretation via cell-state transitions. BCR::ABL1 classification is driven by de-differentiation toward stem-like states 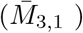. MRD prediction is captured by transitions regulating stem-state persistence and differentiation (including 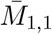 and 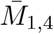).

To understand why the stemness-associated state *x*_1_ was so informative for BCR::ABL1 prediction, we next applied the same machine-learning pipeline to the mechanistically inferred Markov transition rates (Figure 4B; Supplementary Table S2). Among Markov-feature subsets, the best mechanistic feature was 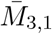, based on its highest composite score (0.81) and minimum BIC (102.66). This result indicates that the de-differentiation transition from state 3 (CD34^−^CD38^+^) back to state 1 (CD34^+^CD38^−^) is not only the most informative mechanistic predictor, but also the transition process that may explain why stemness is the strongest clinical feature for predicting BCR::ABL1-positive (Figure 4C).

For MRD classification, we repeat the same comparison between baseline flow-cytometry features and mechanistically derived Markov features (Figure 4D–F; Supplementary Tables S3–S4). Among flow-cytometry subsets, the best model was again the single-feature model *x*_1_, with composite score 0.67 and BIC 98.49 (Figure 4D; Supplementary Table S3). To understand why the stemness-associated state *x*_1_ was the strongest clinical feature for predicting MRD positivity, we next applied the same machine-learning pipeline to the mechanistically inferred Markov transition rates (Figure 4E; Supplementary Table S4). Among Markov-feature subsets, the primary mechanistic feature was 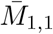,indicating that self-renewal within the stem-like compartment is the dominant transition associated with MRD status. Adding 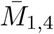, which represents transition out of the stem-like compartment, yielded the best overall Markov model, with the highest composite score (0.71) and minimum BIC (98.58). Together, these results indicate that MRD positivity is associated with the balance between stem-state persistence and differentiation-directed exit, suggesting a mechanistic explanation for why stemness emerges as the strongest clinical predictor in MRD positive (Figure 4F).

Having identified the most informative transitions in Figure 4, we next examined their classification performance and patient-level differences in Figure 5. Figure 5A shows the Receiver Operating Characteristic (ROC) curves for predicting BCR::ABL1 positivity and MRD positivity using the transitions derived from our mechanistic-learning approach. Figure 5B–C then shows how these selected transitions contribute to each endpoint: higher *M*_3,1_ increases the likelihood of BCR::ABL1 positivity, whereas MRD positivity is characterized by higher *M*_1,1_ together with lower *M*_1,4_. Thus, BCR::ABL1-positive disease is associated with stronger de-differentiation back toward the stem-like state, whereas MRD-positive disease is associated with greater stem-like self-renewal and reduced differentiation-directed exit. We further examined these transitions across individual patients. The transition *M*_3,1_ differs significantly between BCR::ABL1-positive and BCR::ABL1-negative groups (Figure 5D), which is also visually apparent in the top row of the patient-level chord diagrams in Figure 5E, where the transition from state 3 back to state 1 is stronger in the BCR::ABL1-positive group. A similar pattern is observed for MRD in the second row of Figure 5E: MRD-positive patients show higher *M*_1,1_ and relatively lower *M*_1,4_ than MRD-negative patients. Figure 5F confirms that these MRD-associated differences are statistically significant.

**Figure 5:**
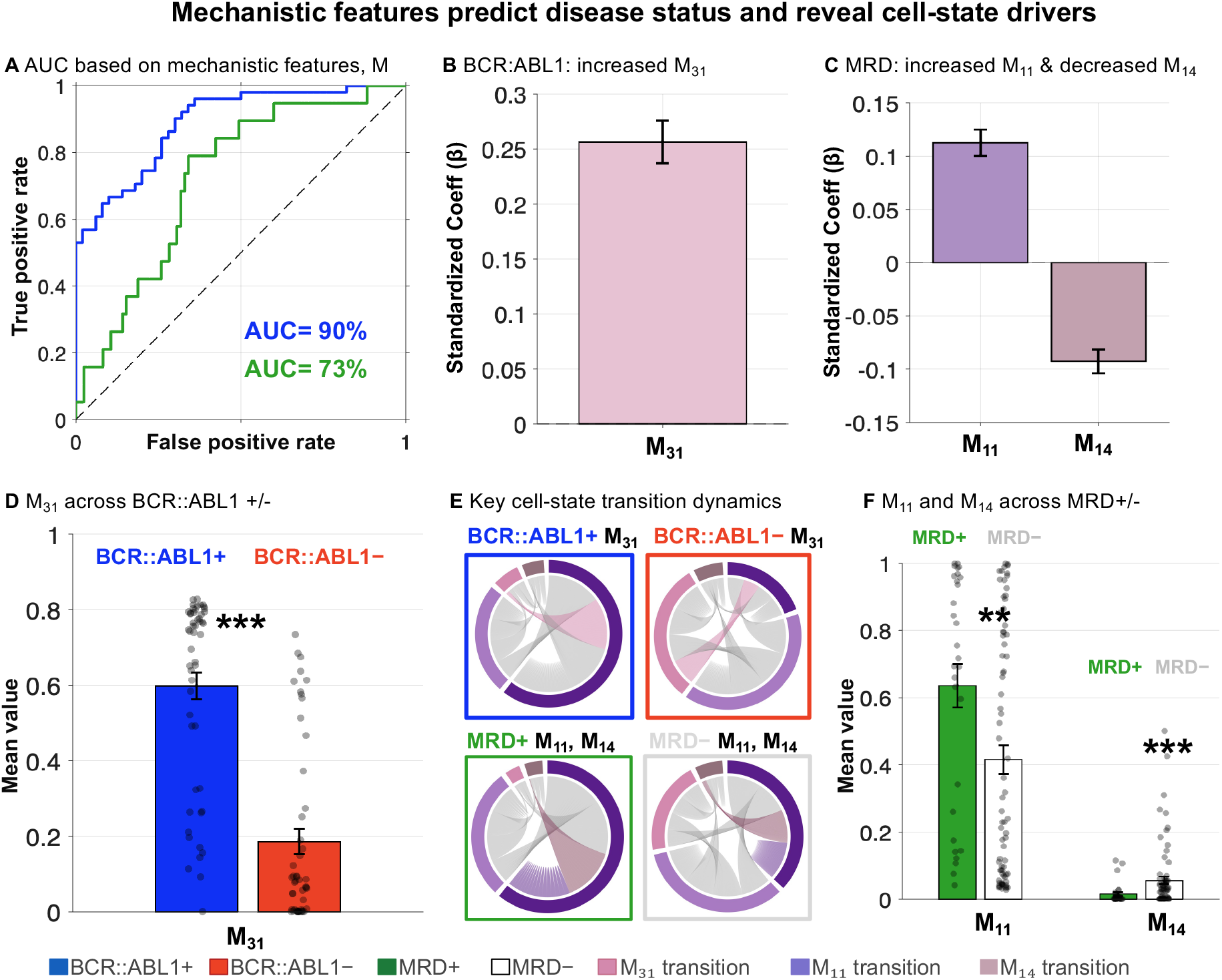
Mechanistic cell-state transitions predict BCR::ABL1 and MRD status and reveal disease-driving dynamics. (A) Receiver operating characteristic curves show that mechanistic features classify BCR::ABL1 status and MRD status with AUCs of 90% and 73%, respectively.(B) The de-differentiation transition *M*_31_ (CD34^−^/CD38^+^ *→* CD34^+^/CD38^−^) is the primary predictor of BCR::ABL1 positivity (*β* ≈ +0.26). (C)MRD status is jointly governed by *M*_11_ self-renewal (*β* ≈ +0.11) and *M*_14_ differentiation (*β* ≈ − 0.09) . (D) The transition *M*_31_ is significantly elevated in BCR::ABL1^+^ patients (mean≈ 0.60 vs 0.19; \*\*\**p* < 0.001). (E) Chord diagrams of patient-level transition matrices confirm qualitatively distinct dynamical landscapes across all four clinical groups. (F) Both *M*_11_ and *M*_14_ significantly distinguish MRD^+^from MRD^−^patients (\*\**p* < 0.01 and \*\*\**p <* 0.001, respectively).

### 3.2 Virtual patients to scale small cohorts attenuates low-dimensional signals

To test whether data augmentation with mechanistically generated virtual patients could improve predictive accuracy and robustness while scaling small cohort sizes, we used the full set of 100 virtual patients generated for each real patient from the fitted Markov model, thereby expanding the cohort 100-fold, and repeated the same weighted ridge-logistic subset-search pipeline used in Figure 1. Figure 6 summarizes the resulting performance for BCR::ABL1 and MRD prediction, and Supplementary Table S5 reports the MRD Markov-feature results in detail.

**Figure 6:**
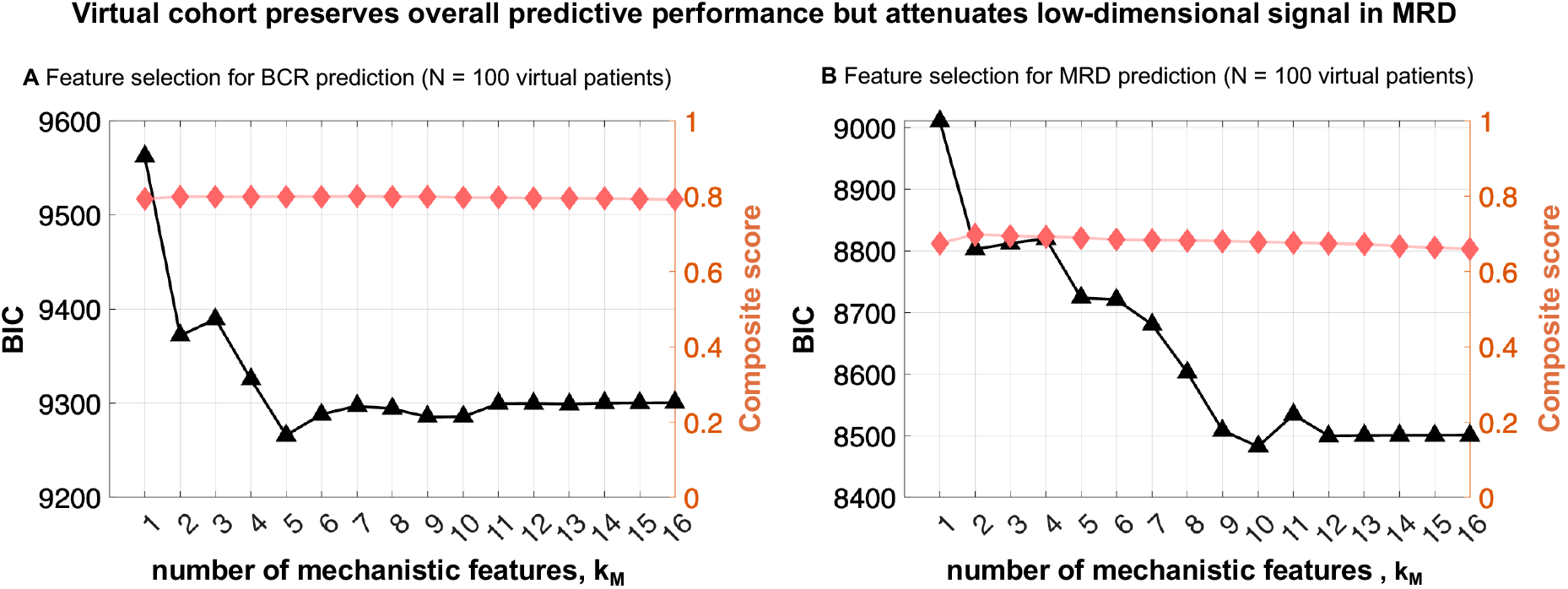
Performance in virtual patient cohort. Feature selection and prediction results for BCR::ABL1 and MRD using virtual patients. Compared to the real cohort, accuracy is reduced across feature types, reflecting increased heterogeneity.

Contrary to our initial expectation, the virtual-cohort analysis did not improve predictive performance. Instead, the composite score was reduced relative to the real cohort, while the minimum BIC shifted toward models with a substantially larger number of features. For MRD, the preferred model in the real cohort was the two-feature subset 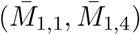 (Figure 4; Supplementary Table S4), whereas in the virtual cohort the best BIC was obtained with the nine-feature model (*M*_1,1_, *M*_1,2_, *M*_1,4_, *M*_2,2_, *M*_2,3_, *M*_2,4_, *M*_3,4_, *M*_4,2_, *M*_4,4_) (Supplementary Table S5). This shift indicates that the augmented cohort does not sharpen the dominant predictive signal, but instead spreads it across a broader set of transition features.

One plausible explanation is that 100-fold expansion amplifies not only the major biological transition patterns, but also rare or weakly estimated transitions. As a result, low-magnitude transition variability is carried forward into the augmented dataset and can dilute the stronger class-separating structure present in the original cohort. These results suggest that virtual patient augmentation is a reasonable method to increase sample size, but it does not automatically improve accuracy or robustness unless the simulated heterogeneity remains tightly aligned with the true underlying biological signal.

### 3.3 Virtual patients can balance subgroups and retain predictive accuracy

We next asked whether virtual patients could be used in place of the class-weighting approach to balance the MRD groups. To address this, we constructed 300 resampled balanced VP datasets, reran the same subset-search pipeline on each dataset (Figure 7), and recorded, for each subset size *k*, the most frequently selected transition combination across runs (Supplementary Table S6). The dominant single transition was *M*_1,1_, and the most frequently selected three-feature subset, (*M*_1,1_, *M*_1,3_, *M*_1,4_), achieved the lowest mean BIC (209.93) with a mean composite score of 0.73. This result is comparable to the MRD model obtained from averaged transitions with class weighting, for which 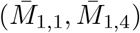 achieved a composite score of 0.71 (Figure 4E; Supplementary Table S4). Thus, VP-based balancing provides only a modest gain in predictive performance, at the cost of increased model complexity and substantially greater computational effort, while yielding a similar biological interpretation centered on stem-like persistence and differentiation-directed exit.

**Figure 7:**
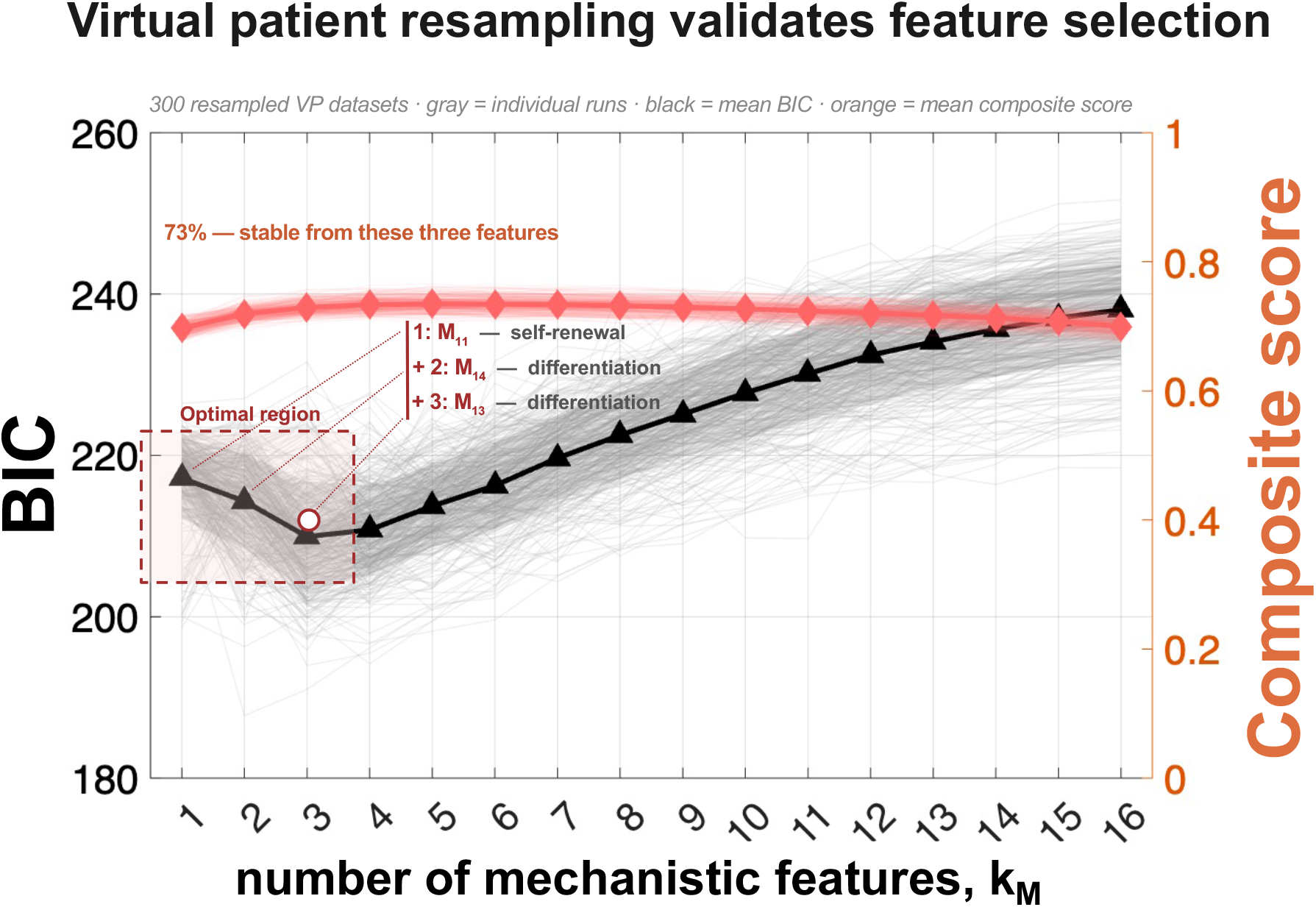
Virtual patient (VP) resampling validates robust feature selection. Model performance was evaluated across 300 balanced VP resampling runs, a number chosen so that each MRD negative VP realization had approximately a 95% probability of being sampled at least once across runs. Gray lines show individual runs, with mean BIC (black) and mean combination score (orange) plotted as a function of the number of mechanistic features. While model fit continues to improve with increasing feature number, predictive performance rapidly saturates, identifying an optimal low-dimensional region (*k* ≈ 2–3). Across resamples, the same minimal feature set, *M*_1,1_ (self-renewal) and *M*_1,4_*/M*_1,3_ (differentiation), is consistently recovered, matching the features identified using class weighting. These results confirm the robustness of mechanistic feature selection.

## 4 Discussion

The goal of this study was to test whether mechanistically derived cell fate transition features can match the predictive utility of standard flow-cytometry features while improving biological interpretability. To do this, we used the same weighted ridge-logistic pipeline for both feature families, ranked feature subsets under the same cross-validation framework, and then examined how the selected transitions appeared in classifier performance and patient-level transition patterns (Figures 1–5). Rather than introducing a new machine learning algorithm in this manuscript, we instead implement a novel pipeline to combine standard mechanistic modeling and machine learning methods, to assess whether mechanistic abstraction improves not only predictive performance, but also the biological interpretability of clinically relevant outcomes.

Our results show that mechanistic features are competitive with baseline clinical features for key endpoints. For BCR::ABL1 prediction, the best clinical single-feature model was *x*_1_, whereas the strongest Markov model was 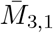(Figure 4A–B; Supplementary Tables S1–S2). For MRD prediction, the preferred flow-cytometry model under the information-criterion ranking was again the single-feature model *x*_1_, whereas the preferred Markov subset was the two-feature model 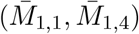 (Figure 4E–F; Supplementary Tables S3–S4). Figure 5 shows that these selected transitions remain visible at the classifier and patient levels, with ROC performance, fitted coefficients, and group-specific transition patterns all supporting the same small set of mechanistic predictors. Taken together, these findings show that a compact set of mechanistic features can preserve predictive signal while pointing directly to the transition processes that drive the biological interpretation discussed next.

An important determinant of the similar accuracy of classifiers trained on biological data features (**X**) versus mechanistic features (**M**) is the small model error. In the initial publication of the mechanistic model described here, Gravenmier et. al. report small error values between the model output and data (eqn. 4) on the order of 10^−5^. This ensures that no information is lost by assessing correlations between biological data and outcomes or assessing correlations between mechanistic parameters and outcome, given the fact that the mechanistic parameters well-describe the data. In the event that a mechanistic model is a poor fit to data, it is expected that our approach may require more mechanistic parameters to maintain the same degree of predictive accuracy as a model trained on the original biological data.

Biologically, the selected transition features are consistent with the mechanistic interpretation established in our prior Markov modeling study[24], but they point to somewhat different aspects of disease biology for BCR::ABL1 and MRD. For BCR::ABL1 classification, the prominence of 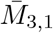 indicates that genotype-associated variation is strongly tied to inward flux from the CD34^−^/CD38^+^compartment back into the stem CD34^+^/CD38^−^state, consistent with the enhanced stem-like reconstruction seen in BCR::ABL1-positive patients in Figure 3. This is also consistent with Figure 2D, where transitions into the stem-like compartment cluster with one another and separate according to BCR::ABL1 status, and with Figure 5B,D–E, where both the fitted coefficient and the patient-level transition diagrams emphasize stronger de-differentiation into the stem compartment. For MRD prediction, repeated selection of 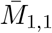 (stem-state persistence) and 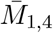 (transition out of the stem compartment) suggests that prognosis-associated variation is driven by how patients balance stem-like persistence versus differentiation-directed exit; Figure 5C,E–F shows that this interpretation is visible both in the classifier coefficients and in the group-specific transition structure. Together, these results suggest that BCR::ABL1 status is most strongly associated with re-entry into the stem-like compartment, whereas MRD is more closely associated with maintenance of that compartment once established.

Finally, the virtual-patient analyses clarify both the usefulness and the limitations of mechanistic augmentation. When the cohort was expanded 100-fold using the full VP set (Figure 6), predictive performance decreased and the minimum-BIC MRD model shifted from the parsimonious two-feature subset 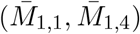 to a substantially larger feature set, suggesting that the simulated heterogeneity spreads predictive signal across weaker transitions rather than sharpening the dominant one. By contrast, when VPs were used specifically to rebalance the MRD classes (Figure 7), the same biological axis centered on stem-state persistence and differentiation-directed exit was repeatedly recovered, and the most frequent three-feature subset (*M*_1,1_, *M*_1,3_, *M*_1,4_) yielded only a modest gain in mean composite score relative to the class-weighted real-cohort model. Together, these results suggest that VP generation is useful as a robustness and balancing tool, but not as a straightforward route to better prediction; in this setting, class weighting captures essentially the same biological signal with lower model complexity and much less computational cost. Overall, these results support mechanistic learning as a practical framework that links predictive modeling to interpretable disease dynamics in B-ALL.

## Acknowledgments

The authors gratefully acknowledge funding by the National Cancer Institute via the Cancer Systems Biology Consortium (CSBC) U54CA274507; and Moffitt Cancer Center support from the Center of Excellence for Evolutionary Therapy.

