## Supplementary Information for "Mechanistic learning to predict and understand minimal residual disease"

### Supplementary Material

TableS1: Flow-cytometry feature subsets to predict BCR::ABL1.

| $k$ | Best Features | Accuracy | BalAcc | AUC | Combo | LogLikelihood | DfEff | BIC |
|---|---|---|---|---|---|---|---|---|
| $\star$ 1 | $x_1$ | 0.76 | 0.76 | 0.88 | 0.80 | -46.29 | 1.93 | 101.47 |
| 2 | $x_1, x_3$ | 0.75 | 0.75 | 0.88 | 0.79 | -45.51 | 2.77 | 103.79 |
| 3 | $x_1, x_2, x_3$ | 0.76 | 0.76 | 0.88 | 0.80 | -45.56 | 3.14 | 105.60 |
| 4 | $x_1$–$x_4$ | 0.74 | 0.74 | 0.88 | 0.79 | -45.18 | 3.81 | 107.95 |

TableS2: Use Markov transition rates to predict BCR::ABL1.

| $k$ | Best Features | Acc | BalAcc | AUC | Combo | LogLikelihood | DfEff | BIC |
|---|---|---|---|---|---|---|---|---|
| $\star$ 1 | $\bar{M}_{31}$ | 0.76 | 0.76 | 0.90 | 0.81 | -46.87 | 1.93 | 102.66 |
| 2 | $\bar{M}_{31}, \bar{M}_{41}$ | 0.76 | 0.76 | 0.89 | 0.81 | -47.06 | 2.00 | 103.35 |
| 3 | $\bar{M}_{13}, \bar{M}_{41}, \bar{M}_{43}$ | 0.78 | 0.78 | 0.86 | 0.81 | -46.71 | 2.85 | 106.59 |
| 4 | $\bar{M}_{13}, \bar{M}_{22}, \bar{M}_{31}, \bar{M}_{33}$ | 0.77 | 0.77 | 0.88 | 0.81 | -46.40 | 3.26 | 107.84 |
| 5 | $\bar{M}_{11}, \bar{M}_{21}, \bar{M}_{31}, \bar{M}_{34}, \bar{M}_{41}$ | 0.76 | 0.76 | 0.89 | 0.80 | -46.44 | 3.11 | 107.25 |
| 6 | $\bar{M}_{11}, \bar{M}_{12}, \bar{M}_{22}, \bar{M}_{23}, \bar{M}_{31}, \bar{M}_{41}$ | 0.76 | 0.76 | 0.89 | 0.80 | -45.45 | 3.52 | 107.13 |
| 7 | $\bar{M}_{13}, \bar{M}_{22}, \bar{M}_{23}, \bar{M}_{31}, \bar{M}_{33}, \bar{M}_{41}, \bar{M}_{44}$ | 0.77 | 0.77 | 0.87 | 0.81 | -46.10 | 4.03 | 110.80 |
| 8 | $\bar{M}_{11}, \bar{M}_{12}, \bar{M}_{21}, \bar{M}_{22}, \bar{M}_{31}, \bar{M}_{32}, \bar{M}_{41}, \bar{M}_{43}$ | 0.76 | 0.76 | 0.88 | 0.80 | -45.45 | 3.75 | 108.20 |
| 9 | $\bar{M}_{13}, \bar{M}_{14}, \bar{M}_{22}, \bar{M}_{23}, \bar{M}_{31}, \bar{M}_{32}, \bar{M}_{33}, \bar{M}_{41}, \bar{M}_{43}$ | 0.76 | 0.76 | 0.88 | 0.80 | -45.82 | 4.16 | 110.84 |
| 10 | $\bar{M}_{12}, \bar{M}_{13}, \bar{M}_{21}, \bar{M}_{22}, \bar{M}_{23}, \bar{M}_{24}, \bar{M}_{31}, \bar{M}_{33}, \bar{M}_{43}, \bar{M}_{44}$ | 0.76 | 0.76 | 0.88 | 0.80 | -45.09 | 4.49 | 110.89 |
| 11 | $\bar{M}_{11}, \bar{M}_{12}, \bar{M}_{21}, \bar{M}_{22}, \bar{M}_{23}, \bar{M}_{24}, \bar{M}_{31}, \bar{M}_{32}, \bar{M}_{34}, \bar{M}_{41}, \bar{M}_{42}$ | 0.76 | 0.76 | 0.87 | 0.80 | -43.96 | 4.65 | 109.36 |
| 12 | $\bar{M}_{11}, \bar{M}_{12}, \bar{M}_{21}, \bar{M}_{22}, \bar{M}_{23}, \bar{M}_{24}, \bar{M}_{31}, \bar{M}_{33}, \bar{M}_{41}, \bar{M}_{42}, \bar{M}_{43}, \bar{M}_{44}$ | 0.75 | 0.75 | 0.88 | 0.80 | -44.65 | 4.62 | 110.63 |
| 13 | $\bar{M}_{11}, \bar{M}_{12}, \bar{M}_{13}, \bar{M}_{14}, \bar{M}_{21}, \bar{M}_{22}, \bar{M}_{23}, \bar{M}_{31}, \bar{M}_{32}, \bar{M}_{33}, \bar{M}_{41}, \bar{M}_{42}, \bar{M}_{43}$ | 0.75 | 0.75 | 0.88 | 0.80 | -44.82 | 4.48 | 110.29 |
| 14 | $\bar{M}_{11}, \bar{M}_{12}, \bar{M}_{13}, \bar{M}_{21}, \bar{M}_{22}, \bar{M}_{23}, \bar{M}_{24}, \bar{M}_{31}, \bar{M}_{32}, \bar{M}_{33}, \bar{M}_{41}, \bar{M}_{42}, \bar{M}_{43}, \bar{M}_{44}$ | 0.75 | 0.75 | 0.88 | 0.79 | -44.62 | 4.73 | 111.05 |
| 15 | $\bar{M}_{11}, \bar{M}_{12}, \bar{M}_{13}, \bar{M}_{21}, \bar{M}_{22}, \bar{M}_{23}, \bar{M}_{24}, \bar{M}_{31}, \bar{M}_{32}, \bar{M}_{33}, \bar{M}_{34}, \bar{M}_{41}, \bar{M}_{42}, \bar{M}_{43}, \bar{M}_{44}$ | 0.74 | 0.74 | 0.88 | 0.79 | -43.91 | 4.99 | 110.83 |
| 16 | $\bar{M}_{11}, \bar{M}_{12}, \bar{M}_{13}, \bar{M}_{14}, \bar{M}_{21}, \bar{M}_{22}, \bar{M}_{23}, \bar{M}_{24}, \bar{M}_{31}, \bar{M}_{32}, \bar{M}_{33}, \bar{M}_{34}, \bar{M}_{41}, \bar{M}_{42}, \bar{M}_{43}, \bar{M}_{44}$ | 0.74 | 0.74 | 0.88 | 0.79 | -43.73 | 5.28 | 111.80 |

TableS3: Use flow cytometry to predict MRD.

| $k$ | Best Features | Accuracy | BalAcc | AUC | Combo | LogLikelihood | DfEff | BIC |
|---|---|---|---|---|---|---|---|---|
| $\star$ 1 | $x_1$ | 0.62 | 0.69 | 0.70 | 0.67 | -44.78 | 1.92 | 98.49 |
| 2 | $x_1, x_4$ | 0.67 | 0.72 | 0.71 | 0.70 | -43.60 | 2.63 | 99.40 |
| 3 | $x_1, x_2, x_4$ | 0.68 | 0.72 | 0.70 | 0.70 | -42.36 | 3.21 | 99.61 |
| 4 | $x_1$–$x_4$ | 0.64 | 0.70 | 0.72 | 0.69 | -41.79 | 3.31 | 98.97 |

TableS4: Use Markov transition rates to predict MRD.

| $k$ | Best Features | Acc | BalAcc | AUC | Combo | LogLikelihood | DfEff | BIC |
|---|---|---|---|---|---|---|---|---|
| 1 | $\bar{M}_{11}$ | 0.64 | 0.70 | 0.72 | 0.69 | -44.83 | 1.92 | 98.60 |
| $\star$ 2 | $\bar{M}_{11}, \bar{M}_{14}$ | 0.67 | 0.72 | 0.73 | 0.71 | -43.29 | 2.58 | 98.58 |
| 3 | $\bar{M}_{11}, \bar{M}_{14}, \bar{M}_{22}$ | 0.67 | 0.72 | 0.72 | 0.70 | -42.43 | 3.17 | 99.58 |
| 4 | $\bar{M}_{11}, \bar{M}_{12}, \bar{M}_{14}, \bar{M}_{34}$ | 0.67 | 0.72 | 0.72 | 0.70 | -41.96 | 3.55 | 100.42 |
| 5 | $\bar{M}_{11}, \bar{M}_{12}, \bar{M}_{13}, \bar{M}_{14}, \bar{M}_{34}$ | 0.66 | 0.71 | 0.73 | 0.70 | -41.65 | 3.69 | 100.41 |
| 6 | $\bar{M}_{11}, \bar{M}_{14}, \bar{M}_{21}, \bar{M}_{23}, \bar{M}_{24}, \bar{M}_{34}$ | 0.66 | 0.71 | 0.72 | 0.70 | -41.51 | 4.10 | 102.06 |
| 7 | $\bar{M}_{11}, \bar{M}_{14}, \bar{M}_{21}, \bar{M}_{22}, \bar{M}_{24}, \bar{M}_{31}, \bar{M}_{44}$ | 0.67 | 0.72 | 0.70 | 0.70 | -41.76 | 4.14 | 102.72 |
| 8 | $\bar{M}_{11}, \bar{M}_{14}, \bar{M}_{21}, \bar{M}_{22}, \bar{M}_{24}, \bar{M}_{31}, \bar{M}_{34}, \bar{M}_{44}$ | 0.67 | 0.72 | 0.70 | 0.70 | -41.69 | 4.35 | 103.59 |
| 9 | $\bar{M}_{11}, \bar{M}_{14}, \bar{M}_{21}, \bar{M}_{22}, \bar{M}_{23}, \bar{M}_{24}, \bar{M}_{34}, \bar{M}_{42}, \bar{M}_{44}$ | 0.66 | 0.71 | 0.72 | 0.70 | -41.37 | 4.47 | 103.51 |
| 10 | $\bar{M}_{11}, \bar{M}_{14}, \bar{M}_{21}, \bar{M}_{22}, \bar{M}_{23},$ $\bar{M}_{24}, \bar{M}_{32}, \bar{M}_{34}, \bar{M}_{42}, \bar{M}_{44}$ | 0.66 | 0.71 | 0.71 | 0.69 | -41.37 | 4.54 | 103.82 |
| 11 | $\bar{M}_{11}, \bar{M}_{12}, \bar{M}_{14}, \bar{M}_{21}, \bar{M}_{22}, \bar{M}_{23},$ $\bar{M}_{24}, \bar{M}_{32}, \bar{M}_{34}, \bar{M}_{42}, \bar{M}_{44}$ | 0.65 | 0.70 | 0.71 | 0.69 | -41.33 | 4.63 | 104.18 |
| 12 | $\bar{M}_{11}, \bar{M}_{12}, \bar{M}_{14}, \bar{M}_{21}, \bar{M}_{22}, \bar{M}_{23},$ $\bar{M}_{24}, \bar{M}_{31}, \bar{M}_{33}, \bar{M}_{34}, \bar{M}_{41}, \bar{M}_{44}$ | 0.64 | 0.70 | 0.71 | 0.68 | -41.30 | 4.63 | 104.12 |
| 13 | $\bar{M}_{11}, \bar{M}_{13}, \bar{M}_{14}, \bar{M}_{21}, \bar{M}_{22}, \bar{M}_{23}, \bar{M}_{24},$ $\bar{M}_{31}, \bar{M}_{34}, \bar{M}_{41}, \bar{M}_{42}, \bar{M}_{43}, \bar{M}_{44}$ | 0.64 | 0.70 | 0.71 | 0.68 | -41.27 | 4.63 | 104.05 |
| 14 | $\bar{M}_{11}, \bar{M}_{12}, \bar{M}_{14}, \bar{M}_{21}, \bar{M}_{22}, \bar{M}_{23}, \bar{M}_{24},$ $\bar{M}_{31}, \bar{M}_{33}, \bar{M}_{34}, \bar{M}_{41}, \bar{M}_{42}, \bar{M}_{43}, \bar{M}_{44}$ | 0.64 | 0.70 | 0.71 | 0.68 | -41.26 | 4.71 | 104.41 |
| 15 | $\bar{M}_{11}, \bar{M}_{12}, \bar{M}_{13}, \bar{M}_{14}, \bar{M}_{21}, \bar{M}_{22}, \bar{M}_{23}, \bar{M}_{24},$ $\bar{M}_{31}, \bar{M}_{33}, \bar{M}_{34}, \bar{M}_{41}, \bar{M}_{42}, \bar{M}_{43}, \bar{M}_{44}$ | 0.63 | 0.69 | 0.71 | 0.68 | -41.24 | 4.74 | 104.51 |
| 16 | $\bar{M}_{11}, \bar{M}_{12}, \bar{M}_{13}, \bar{M}_{14}, \bar{M}_{21}, \bar{M}_{22}, \bar{M}_{23}, \bar{M}_{24},$ $\bar{M}_{31}, \bar{M}_{32}, \bar{M}_{33}, \bar{M}_{34}, \bar{M}_{41}, \bar{M}_{42}, \bar{M}_{43}, \bar{M}_{44}$ | 0.64 | 0.70 | 0.71 | 0.68 | -41.23 | 4.79 | 104.73 |

TableS5: Use Markov transition rates of virtual patients to predict MRD.

| $k$ | Best Features | Acc | BalAcc | AUC | Combo | LogLikelihood | DfEff | BIC |
|---|---|---|---|---|---|---|---|---|
| 1 | $M_{11}$ | 0.63 | 0.68 | 0.72 | 0.67 | -4496.29 | 2.00 | 9011.07 |
| 2 | $M_{11}, M_{14}$ | 0.66 | 0.71 | 0.73 | 0.70 | -4387.63 | 2.99 | 8802.95 |
| 3 | $M_{11}, M_{14}, M_{44}$ | 0.66 | 0.71 | 0.72 | 0.70 | -4387.63 | 3.99 | 8812.18 |
| 4 | $M_{11}, M_{14}, M_{22}, M_{24}$ | 0.67 | 0.71 | 0.71 | 0.70 | -4335.76 | 4.99 | 8717.65 |
| 5 | $M_{11}, M_{14}, M_{22}, M_{34}, M_{44}$ | 0.67 | 0.71 | 0.71 | 0.70 | -4336.43 | 5.99 | 8728.23 |
| 6 | $M_{11}, M_{14}, M_{22}, M_{24}, M_{34}, M_{44}$ | 0.67 | 0.71 | 0.71 | 0.70 | -4335.31 | 6.98 | 8735.23 |
| 7 | $M_{11}, M_{14}, M_{22}, M_{24}, M_{34}, M_{42}, M_{44}$ | 0.67 | 0.72 | 0.70 | 0.70 | -4323.42 | 7.98 | 8720.66 |
| 8 | $M_{11}, M_{14}, M_{22}, M_{24}, M_{32}, M_{34}, M_{42}, M_{44}$ | 0.67 | 0.71 | 0.70 | 0.70 | -4320.41 | 8.98 | 8723.84 |
| $\star$ 9 | $M_{11}, M_{12}, M_{14}, M_{22}, M_{23}, M_{24}, M_{34}, M_{42}, M_{44}$ | 0.65 | 0.71 | 0.72 | 0.69 | -4191.38 | 9.88 | 8474.15 |
| 10 | $M_{11}, M_{12}, M_{14}, M_{22}, M_{23},$ $M_{24}, M_{32}, M_{34}, M_{42}, M_{44}$ | 0.65 | 0.70 | 0.72 | 0.69 | -4191.16 | 10.88 | 8482.93 |
| 11 | $M_{11}, M_{12}, M_{14}, M_{21}, M_{22}, M_{23},$ $M_{24}, M_{32}, M_{34}, M_{42}, M_{44}$ | 0.65 | 0.70 | 0.72 | 0.69 | -4191.13 | 10.88 | 8482.95 |
| 12 | $M_{11}, M_{12}, M_{14}, M_{21}, M_{22}, M_{23},$ $M_{24}, M_{31}, M_{32}, M_{34}, M_{42}, M_{44}$ | 0.64 | 0.70 | 0.72 | 0.69 | -4190.79 | 11.86 | 8491.30 |
| 13 | $M_{11}, M_{12}, M_{14}, M_{21}, M_{22}, M_{23}, M_{24},$ $M_{31}, M_{32}, M_{34}, M_{41}, M_{42}, M_{44}$ | 0.64 | 0.70 | 0.72 | 0.69 | -4190.76 | 12.84 | 8500.27 |
| 14 | $M_{11}, M_{12}, M_{14}, M_{21}, M_{22}, M_{23}, M_{24},$ $M_{31}, M_{32}, M_{34}, M_{41}, M_{42}, M_{43}, M_{44}$ | 0.63 | 0.69 | 0.72 | 0.68 | -4190.76 | 12.85 | 8500.42 |
| 15 | $M_{11}, M_{13}, M_{14}, M_{21}, M_{22}, M_{23}, M_{24},$ $M_{31}, M_{32}, M_{33}, M_{34}, M_{41}, M_{42}, M_{43}, M_{44}$ | 0.63 | 0.69 | 0.72 | 0.68 | -4190.58 | 12.92 | 8500.66 |
| 16 | $M_{11}, M_{12}, M_{13}, M_{14}, M_{21}, M_{22}, M_{23}, M_{24},$ $M_{31}, M_{32}, M_{33}, M_{34}, M_{41}, M_{42}, M_{43}, M_{44}$ | 0.63 | 0.69 | 0.72 | 0.68 | -4190.58 | 12.93 | 8500.79 |

TableS6: MRD prediction summary across 300 VP-resampling runs used to balance the data.

| $k$ | Most Frequent Combination | Count | % Iterations | Mean BIC | Mean Acc | Mean BalAcc | Mean AUC | Mean Combo |
|---|---|---|---|---|---|---|---|---|
| 1 | $M_{11}$ | 174 | 58.00 | 217.17 | 0.69 | 0.69 | 0.72 | 0.70 |
| 2 | $M_{11}, M_{14}$ | 118 | 39.33 | 214.33 | 0.71 | 0.71 | 0.73 | 0.72 |
| $\star$ 3 | $M_{11}, M_{13}, M_{14}$ | 26 | 8.67 | 209.93 | 0.72 | 0.72 | 0.74 | 0.73 |
| 4 | $M_{11}, M_{13}, M_{14}, M_{23}$ | 10 | 3.33 | 210.78 | 0.73 | 0.73 | 0.75 | 0.73 |
| 5 | $M_{13}, M_{14}, M_{21}, M_{34}, M_{44}$ | 5 | 1.67 | 213.72 | 0.73 | 0.73 | 0.75 | 0.73 |
| 6 | $M_{11}, M_{12}, M_{13}, M_{14}, M_{22}, M_{24}$ | 3 | 1.00 | 216.27 | 0.73 | 0.73 | 0.75 | 0.73 |
| 7 | $M_{11}, M_{12}, M_{13}, M_{14}, M_{22}, M_{24}, M_{42}$ | 3 | 1.00 | 219.67 | 0.73 | 0.73 | 0.74 | 0.73 |
| 8 | $M_{11}, M_{13}, M_{14}, M_{23}, M_{24}, M_{34}, M_{43}, M_{44}$ | 4 | 1.33 | 222.51 | 0.73 | 0.73 | 0.74 | 0.73 |
| 9 | $M_{11}, M_{12}, M_{13}, M_{14}, M_{22}, M_{24}, M_{32}, M_{42}, M_{44}$ | 4 | 1.33 | 225.10 | 0.73 | 0.73 | 0.74 | 0.73 |
| 10 | $M_{12}, M_{13}, M_{14}, M_{21}, M_{22}, M_{23},$ $M_{24}, M_{34}, M_{43}, M_{44}$ | 3 | 1.00 | 227.75 | 0.72 | 0.72 | 0.74 | 0.73 |
| 11 | $M_{11}, M_{12}, M_{13}, M_{14}, M_{22}, M_{23}, M_{24},$ $M_{32}, M_{34}, M_{42}, M_{44}$ | 3 | 1.00 | 230.12 | 0.72 | 0.72 | 0.73 | 0.72 |
| 12 | $M_{11}, M_{12}, M_{13}, M_{14}, M_{21}, M_{22}, M_{23}, M_{24},$ $M_{33}, M_{34}, M_{43}, M_{44}$ | 4 | 1.33 | 232.44 | 0.72 | 0.72 | 0.73 | 0.72 |
| 13 | $M_{11}, M_{12}, M_{13}, M_{14}, M_{21}, M_{22}, M_{23}, M_{24},$ $M_{32}, M_{34}, M_{41}, M_{42}, M_{44}$ | 9 | 3.00 | 234.08 | 0.71 | 0.71 | 0.73 | 0.72 |
| 14 | $M_{11}, M_{12}, M_{13}, M_{14}, M_{21}, M_{22}, M_{23}, M_{24},$ $M_{32}, M_{34}, M_{41}, M_{42}, M_{43}, M_{44}$ | 16 | 5.33 | 235.64 | 0.71 | 0.71 | 0.73 | 0.71 |
| 15 | $M_{11}, M_{12}, M_{13}, M_{14}, M_{21}, M_{22}, M_{23}, M_{24},$ $M_{32}, M_{33}, M_{34}, M_{41}, M_{42}, M_{43}, M_{44}$ | 37 | 12.33 | 237.04 | 0.70 | 0.70 | 0.72 | 0.71 |
| 16 | $M_{11}, M_{12}, M_{13}, M_{14}, M_{21}, M_{22}, M_{23}, M_{24},$ $M_{31}, M_{32}, M_{33}, M_{34}, M_{41}, M_{42}, M_{43}, M_{44}$ | 300 | 100.00 | 238.07 | 0.69 | 0.69 | 0.72 | 0.70 |
